# Blood Unit Segments Accurately Represent the Biophysical Properties of Red Blood Cells in Blood Bags but Not Hemolysis

**DOI:** 10.1101/2021.07.20.453156

**Authors:** Emel Islamzada, Kerryn Matthews, Erik Lamoureux, Simon P. Duffy, Mark D. Scott, Hongshen Ma

## Abstract

**BACKGROUND:** The biophysical properties of red blood cells (RBCs) provide potential biomarkers for the quality of donated blood. Blood unit segments provide a simple and nondestructive way to sample RBCs in clinical studies of transfusion efficacy, but it is not known whether RBCs sampled from segments accurately represent the biophysical properties of RBCs in blood bags.

**STUDY DESIGN AND METHODS:** RBCs were sampled from blood bags and segments every two weeks during 8 weeks of storage at 4 °C. RBC deformability was measured by deformability-based sorting using the microfluidic ratchet device in order to derive a rigidity score. Standard hematological parameters, including mean corpuscular volume (MCV), red cell distribution width (RDW), mean cell hemoglobin (MCH), mean corpuscular hemoglobin concentration (MCHC), and hemolysis were measured at the same time points.

**RESULTS:** Deformability of RBCs stored in blood bags was retained over 4 weeks storage but a progressive loss of deformability was observed at weeks 6 and 8. This trend was mirrored in blood unit segments with a strong correlation to the blood bag data. Strong correlations were also observed between blood bag and segment for MCV, MCHC and MCH, but not for hemolysis.

**CONCLUSION:** RBCs sampled from blood unit segments accurately represents the biophysical properties of RBCs in blood bags, but not hemolysis. Blood unit segments provide a simple and non-destructive sample for measuring RBC biophysical properties in clinical studies.

## INTRODUCTION

Red blood cell (RBC) transfusion is a potentially life-saving procedure to restore tissue oxygenation for patients suffering from acute and chronic anemia. Storage of donated RBC is currently limited to 42 days because of the development of the *storage lesion*, which include a series of structural (lipid peroxidation, Band 3 aggregation, membrane asymmetry), metabolic (slowed metabolism due to ATP and 2,3-diphosphoglycerate depletion), and morphologic transformations (discoid, echinocyte, and spherocyte)^1–4^. Development of the storage lesion results in increased RBC hemolysis pre-transfusion and accelerated clearance of transfused RBCs posttransfusion, both of which reduce the clinical benefit of the transfused RBCs^5^. Accelerated RBC clearance is especially problematic for chronic transfusion recipients, who will then require more frequent transfusions and be at greater risk of morbidities, such as iron overload^6^ and transfusion related acute lung injury (TRALI) ^7–10^.

A property common to all of the changes associated with the storage lesion is the reduction of RBC deformability^3,11–14^. This consistent characteristic of RBC storage could therefore serve as a biomarker for the storage lesion. Furthermore, loss of RBC deformability may correspond to post-transfusion efficacy since reduced deformability impedes the transit of these cells through the microvasculature and because more rigid RBCs are retained by the reticuloendothelial system and subsequently eliminated by macrophages^5,15^. As a result, less deformable RBCs have dramatically reduced circulation time and therapeutic efficacy^16,17^. RBCs preserved in cold storage demonstrate an apparent accelerated senescence, characterized by a significant decrease in RBC deformability within three weeks^3,11,18–21^. The more rigid RBCs exhibit a dramatically reduced circulation time, from 120 days to, in some cases, hours or minutes^22–30^. This accelerated senescence corresponds with an increased frequency of post-surgery mortality, following transfusion of blood stored more than two weeks^31^. Importantly, however, we and others have reported that RBC deformability loss is donor dependent, where some donors are able to provide highly stable RBCs while other donors provide rapidly degrading RBCs^2,3,32–34^.

In order to perform clinical studies of how donor RBCs affect transfusion outcomes, it is necessary to sample donor RBCs transfused into recipients. However, sampling from donor blood bags at the point of transfusion is very difficult in practice because of sterility requirements, limited time-window available for transfusions, as well as a general reluctance to deviate from an established protocol with an excellent safety profile. A compelling alternative approach is to sample RBCs from blood bag segments instead of the blood bag. Blood bag segments are blood-filled tubing that remain attached to the blood bag and are typically heat-sealed at several places to allow for serological and hemodynamic assay without compromising the blood unit. Assessing RBC quality from segments is challenging because standard parameters, such as hematocrit, hemoglobin content, and percent hemolysis, have been shown to diverge significantly between segment and blood bag by day 5 of storage^35^. However, changes in RBC deformability represent the cumulative effect of many cellular transformations and may constitute a more robust biomarker for blood quality. To date, it has not been established whether time-dependent changes in RBC deformability are consistent between segment and blood bag during cold storage.

Here, we performed a comparative study of RBC deformability loss when the sample is stored in blood bags and blood bag segments. Using a sensitive microfluidic technique to perform deformability measurements on large numbers of single RBCs, we observed distinct donor-specific RBC aging curves, which demonstrate that the loss of RBC deformability proceeded at different rates for each donor. Importantly, the aging curves obtained from both segment and blood bags were well-matched. Consequently, RBCs sampled from the blood bag segments provide a reliable and non-destructive way to assess the biophysical degradation of RBCs stored in blood bags.

## METHODS

### Packed RBC bags and segments

This study was approved by University of British Columbia’s Clinical Research Ethics Board (UBC REB# H19-01121) and Canadian Blood Services Research Ethics Board (CBS REB# 2019-029). RBC bag units (n=14) were collected and processed by Canadian Blood Services between January 2020 and February 2021. Tubing attached to blood bags were segmented into 4 - 8 sections according to standard practice. RBC units with attached segments were stored according to Canadian Blood Services (CBS) standard operating procedures at 4 °C for 8 weeks.

### RBC sampling and testing

RBCs were sampled from the blood bags and segments to measure their deformability profile, as well as standard hematological parameters and the rate of hemolysis. RBCs are sampled on day 1 of collection (also referred as the week 0 data point), and then at 2, 4, 6 and 8 weeks of cold storage. At the time of testing, 3.0 ml of packed RBCs was drawn through the blood administration port from each bag using a 27-gauge needle (BD). The collection site was then sealed off with Parafilm to maintain sterility throughout storage. RBCs from segments were also collected using a 27-gauge needle, by piercing through the tubing close to the sealed edges. At least 0.4 ml of RBCs was drawn from each segment. In units that did not have enough segments for sampling, priority was made to sample at day 1 (week 0), then week 6, 8 and finally weeks 2 and 4.

At each time point, a 100 μl aliquot of both bag and segment RBCs was removed for monitoring hematological parameters changes that occur during storage. Mean corpuscular volume (MCV), RBC distribution width (RDW), mean cell hemoglobin (MCH), mean corpuscular hemoglobin concentration (MCHC) and hematocrit were measured using the Sysmex Hematology Analyzer. The remaining samples were then centrifuged at 1500 G for 10 minutes to remove the supernatant. Supernatant was stored at −80°C for hemolysis analysis at the end of the study. Packed RBC samples were washed three times in Hank’s Balanced Salt Solution (HBSS, Gibco) with 0.2% Pluronic (F127, MilliporeSigma) at 300 G for 10 minutes. The samples were resuspended at 1% hematocrit in HBSS/0.2% Pluronic prior to deformability measurement.

### RBC deformability analysis

The microfluidic ratchet device sorts RBCs based on their ability to deform through a constriction matrix of different sizes into 12 distinct outlets ^34,36,37^. The RBCs are flowed through the device using an oscillatory (upward/downward) sorting pressure and constant forward pressure system, which moves the sample forward, through the sorting matrix. After sorting the RBCs through the constriction matrix, the number of RBCs in each outlet is counted to obtain a distribution (**Fig. 1A**). Using the distribution, a cumulative frequency can be calculated from which a rigidity score (RS) can be inferred as where the cumulative frequency crosses 50% (**Fig. 1B**), the outlet at which 50% of the sample is collected. The linear interpolation of the cumulative frequency graph can be used to calculate fractional outlet numbers. The RS can be used to compare distributions between different samples as well as different donors (**Fig. 1C**). A loss in deformability is measured as a change in RS throughout storage.

**Fig. 1.**
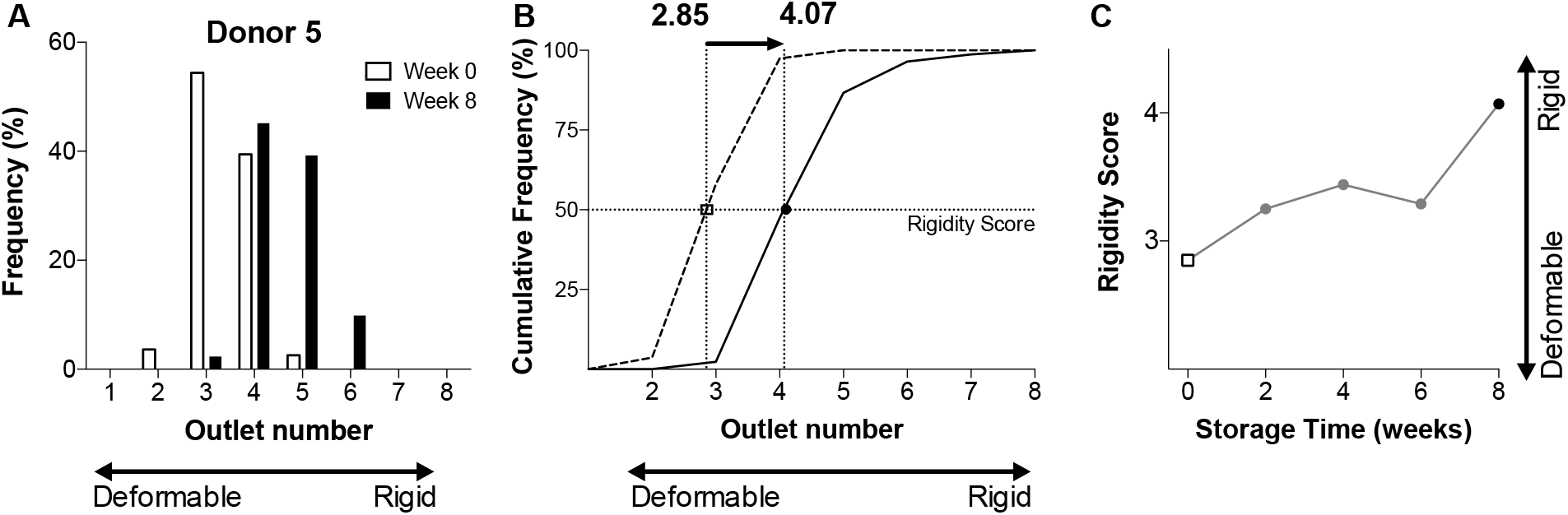
Sample of data analysis. (A) RBCs in each outlet fraction are counted to determine the frequency distribution for each sample. (B) The frequency distribution is converted to cumulative distribution to easily compare samples and donors. A Rigidity Score can be determined from the outlet number where the cumulative frequency distribution crosses 50%. Fractional outlet numbers are determined by linear interpolation between nearest data points. (C) Rigidity scores can be used to track the loss of RBC deformability during storage.

The microfluidic ratchet device was manufactured as previously described^34,36,37^ using photolithographic microfabrication and replica molding of Polydimethylsiloxane (PDMS, Sylgard-184, Ellsworth Adhesives). To measure RBC deformability, the device is first primed by filling with HBSS+0.2% Pluronic buffer for 15 minutes until fully buffered. The microfluidic ratchet device operates using an oscillatory (upward/downward) sorting pressure and constant forward pressure system, which moves the sample forward, through the sorting matrix, towards one of 12 distinct outlets based on the deformability of each RBC. The upward pressure is applied at 175 mbar for 4 seconds. The downward pressure is applied for 1 second at 162 mbar. The forward and sample pressures are applied at 40-45 and 50-55 mbar respectively. The distribution of cells in each distinct outlet is counted after the sorting procedure.

### Hemolysis analysis

Frozen supernatant from 10 matched donor bag and segment samples were fully thawed. An aliquot of each supernatant sample was transferred to a fresh tube and centrifuged at 3000 g for 3 minutes. Packed, unwashed blood samples from day 1 of storage were vortexed for 30 seconds, and diluted 1:10 with DI water. Finally, 10 μl from each sample was transferred to a 96-well flat bottom plate together with 100ul of Drabkin’s reagent (MilliporeSigma) containing Brij-35 solution (Thermofisher). The plate was incubated at room temperature for 15 min on a plate shaker, and absorbance at 450 nm was read using a SpectraMAX Plus plate reader. The amount of hemolysis was then calculated using the following formula:

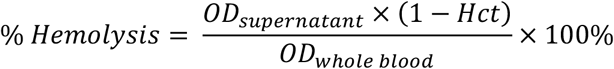

where *Hct* is the hematocrit, and *OD_supernatant_* and *OD_whole blood_* are the measured optical density from the supernatant and whole blood respectively.

### Statistical analysis

Statistical analysis was performed using GraphPad Prism (V8.0) software. Means and standard deviations (SD) are plotted unless otherwise stated. The D’Agostino and Pearson omnibus test was used to determine the normality of all data. Wilcoxon matched pairs signed rank test was used for comparison of data sets. Correlations between normally distributed datasets was calculated using Pearson r with 95% confidence interval.

## RESULTS

### RBC deformability

We established a rigidity score (RS) to measure the deformability of RBCs sampled from blood bags and segments from 14 donors (6 female, 8 male). RBCs were sampled on day 1 (week 0) and on each subsequent 14 days until day 56 (week 8). This sampling period was selected to assess blood bags over the entire 42-day storage window as well as 14 days after outdating. In general, donor RBCs exhibited a progressive deformability loss over time (**Fig. 2**). This deformability loss was more rapid for some donors (e.g. Donor M04 and M05), while other donors demonstrated little-to-no changes in cell deformability (e.g. Donor F03 and F04). Together, these results suggest that monitoring donor blood bags could be used to identify blood bags with a lower apparent RBC deterioration associated with the storage lesion.

**Fig. 2.**
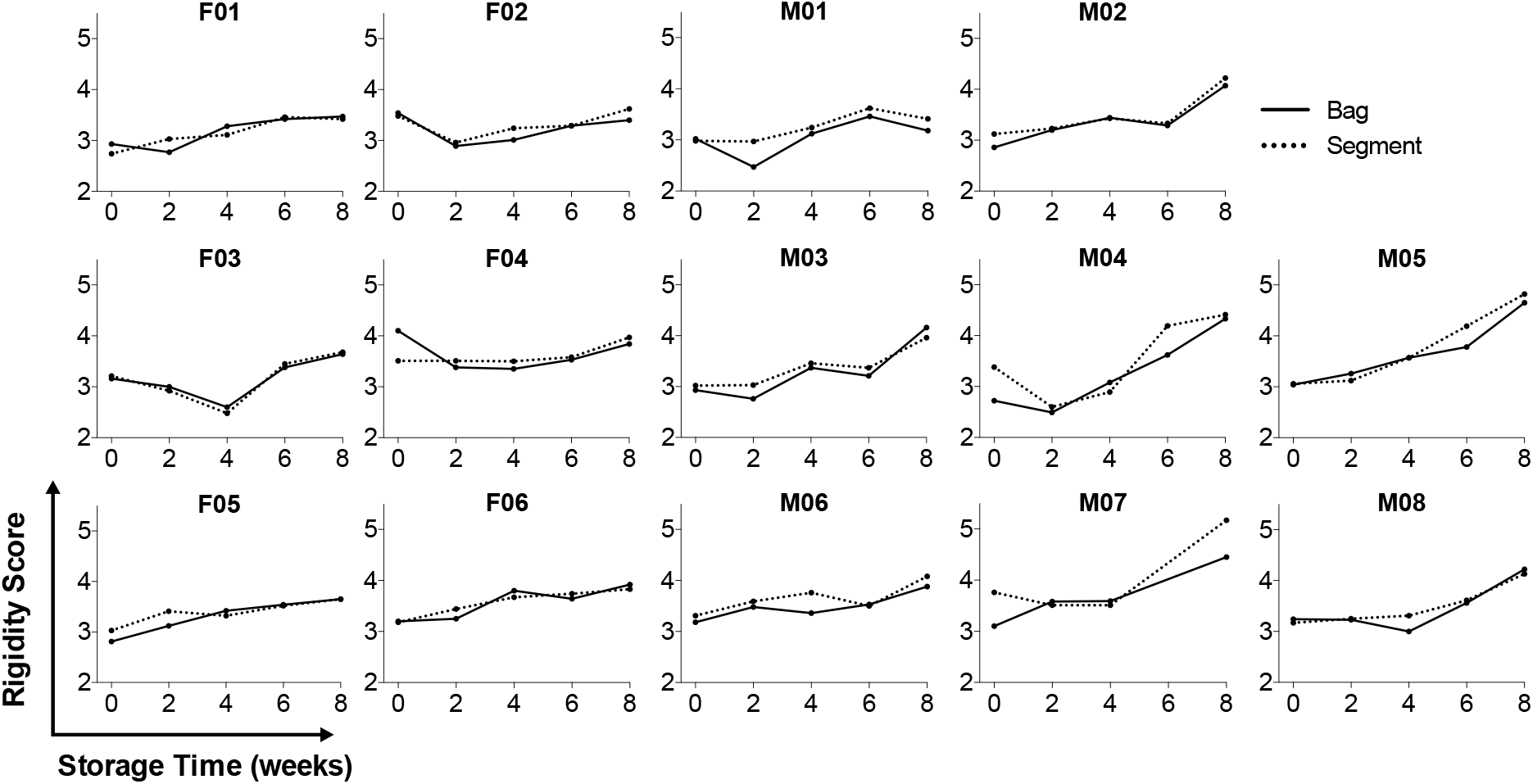
Deformability profiles of RBCs from blood bags and segments during storage. Rigidity Scores of RBCs measured from 14 donors (6 female and 8 male) during 8 weeks of storage in blood bags (solid lines) and their matching segments (dashed lines). F indicates female donors. M indicates male donors.

To investigate sampling of the segment as a non-destructive assessment of RBC deformability in blood bags, we compared the deformability profiles of stored RBCs in each blood bag and corresponding segment (**Fig. 2**). Despite significant inter-donor variability in loss of RBC deformability during storage, we observed direct concordance between the deformability profiles of RBCs stored in blood bags and segment. The mean RS for all donors showed a consistent trend in RBC deformability. Specifically, while no significant change in RBC deformability was observed in weeks 0-4, loss of RBC deformability was observed for both blood bags and segment at week 6 (bag p=0.0164, segment p=0.0010). Loss of RBC deformability was even more pronounced at 8 weeks of storage (bag p=0.0009, segment p=0.0001, **Fig. 2, 3A**). To determine whether deformability of segment RBCs was directly correlated with RBCs from the corresponding blood bag, we performed a linear correlation of RS-values from each (**Fig. 3B**). The correlation between RS-value for blood bags and corresponding segments were significant (Pearson’s r = 0.8863), suggesting that measuring RBC deformability in segment constitutes an accurate and non-destructive measure of the deformability of RBC from the corresponding blood bag, during cold storage.

**Fig. 3.**
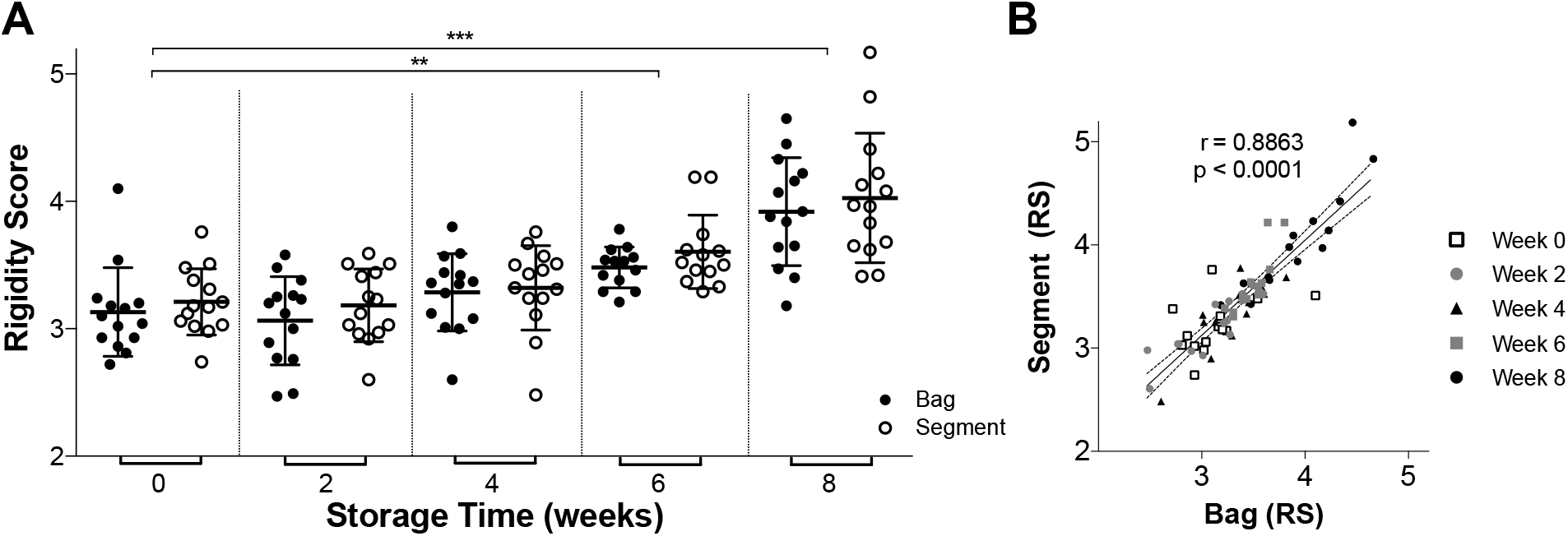
Summary of RBC Rigidity Score during storage. (A) Rigidity Scores for RBCs obtained from blood bags and segments during storage showing detectable deformability loss by 6 weeks (bag p=0.0164, segment p=0.0010), and significant deformability loss by 8 weeks (bag p=0.0009, segment p=0.0001). (B) Correlation between Rigidity Score obtained from RBCs from blood bags and RBCs from matching segments (r = 0.8863).

### Hematological parameters

At each of the timepoints of cold storage, we measured standard hematological parameters on RBCs from bag and segments, including MCV, MCHC, MCH, and RDW using the Sysmex system. There was a strong correlation between RBCs from bag and segments for MCV (r = 0.9419), MCHC (r = 0.9039) and MCH (r = 0.9878). The correlation for RDW was weaker (r = 0.6491), but this is likely a result of the limited dynamic range of the RDW measurement (**Fig. 4**).

**Fig. 4.**
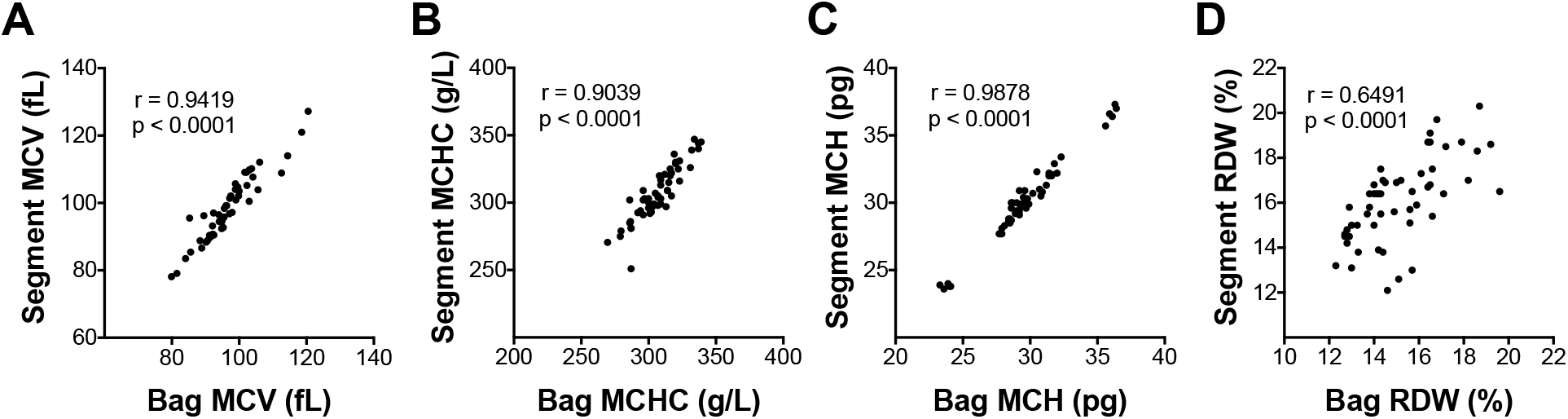
Correlations of RBC hematological parameters between blood bags and segments. (A) MCV (r = 0.9419), (B) MCHC (r = 0.9039), (C) MCH (r = 0.9878), and (D) RDW (r = 0.6491).

### Hemolysis in bags and segments

Hemolysis in bags and segments were measured at each time point by sampling supernatants from the bag and segment. In blood bags, hemolysis levels generally remained below the acceptable threshold of 0.8% from week 0-6. By week 8, hemolysis levels in half of the bags were found to be above the 0.8% threshold (**Fig. 5A**). In blood unit segments, hemolysis was considerably higher. In fact, on the day of production, a majority (6/10) of segments already showed hemolysis levels greater than 0.8%. At four weeks and onwards, the level of hemolysis increased dramatically reaching ~2% by week 6 and ~3.5% by week 8 (**Fig. 5A**). Hemolysis data were also uncorrelated to RS for both bags and segments, which confirmed that RS was independent of hemolysis (**Fig. 5B**).

**Fig. 5.**
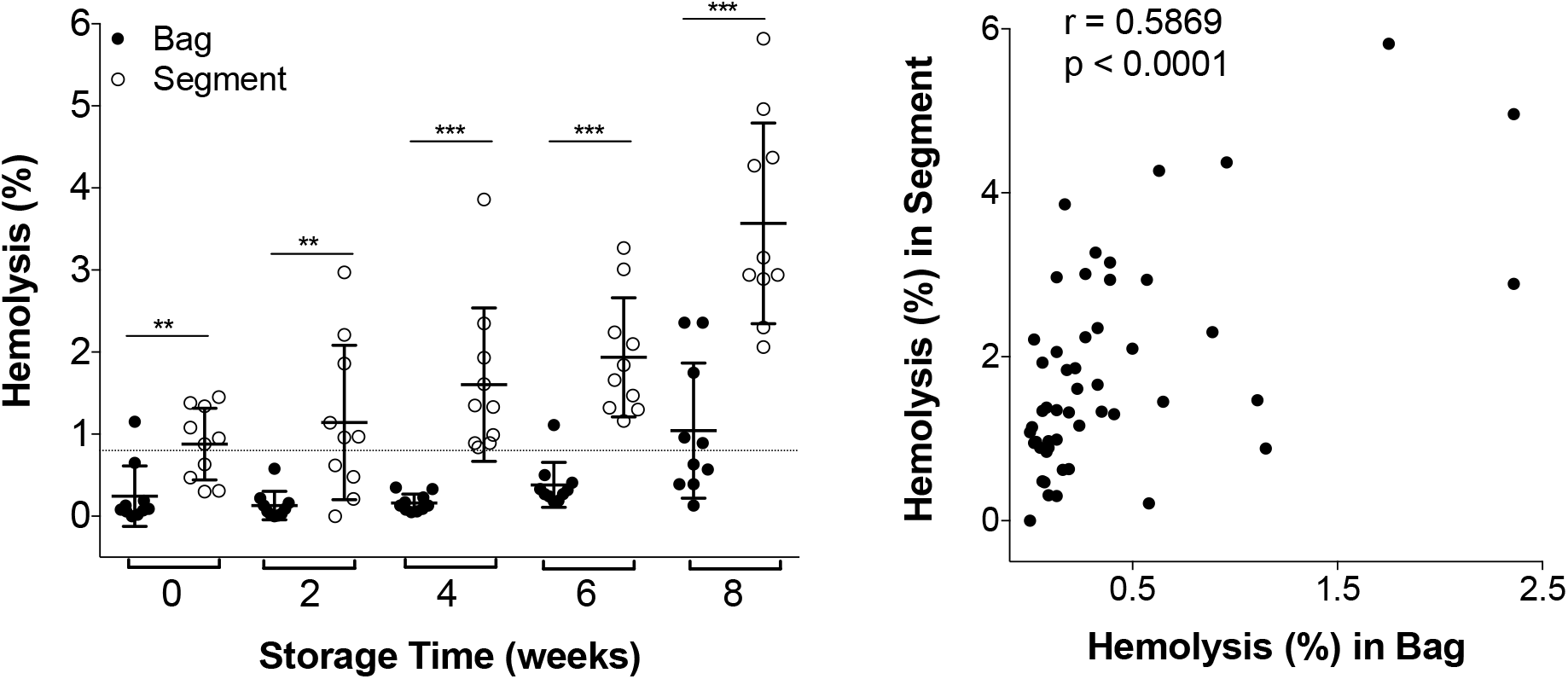
Hemolysis in blood bags and segments. (A) Comparing hemolysis in blood bags (closed circles) and segments (open circles). ** week 0 p=0.0021, ** week 2 p=0.0029, ***week 4, 6 and 8 p<0.0001. (B) Correlation between hemolysis in blood bags and segments.

## DISCUSSION

In this study, we demonstrate that blood unit segments can be used to assess RBC deformability in blood bags without compromising their integrity. This analysis was made using the microfluidic ratchet device to compare RBC deformability in cells from blood bags and segments. As described previously^34^, stored RBCs exhibited a progressive loss of cell deformability. However, individual donor RBCs differed significantly with respect to the magnitude of deformability loss. This variability suggests the potential value in monitoring RBC deterioration in individual blood bags. However, sampling RBCs directly from the blood bag could contaminate and compromise the RBC unit. To address this limitation, we assessed whether the loss of RBC deformability in the blood bag can be inferred by sampling blood unit tubing segments and we found significant correlation between RBC deformability in both blood bag and segment across multiple donors and over the course of eight weeks cold storage. These results demonstrate that progressive loss of RBC deformability can be reliably assessed during blood bag storage by sampling cells from blood unit segments without compromising the blood bag.

By suspending RBCs in preservative solution and storing them in blood bags at 4 °C, RBC units can be stored for up to 42 days prior to transfusion^38,39^. However, stored RBCs undergo a series of structural, metabolic and morphological changes during storage^1–4^ that coincide with reduced transfusion efficacy of stored blood^40–44^. The loss of RBC deformability is a potential biomarker for RBC deterioration during cold storage because these cellular changes collectively contribute to a progressive loss in RBC deformability^3,11–14^. Furthermore, the loss of RBC deformability may facilitate uptake of transfused RBC by reticuloendothelial macrophages^39^ and can predict transfusion outcomes^45^. Our data support previous reports that the rate of RBC degradation differs between donors^2,3,32,33^, suggesting that monitoring RBC deformability during storage can guide blood unit selection to optimize transfusion efficacy for sensitive recipients.

The primary challenge in routine monitoring of RBC deformability in blood bags is that sampling cells can potentially lead to contaminating the RBC unit. The tubing leading into the blood bag provides a compelling source of RBCs, as the tubing can be sealed into segments to enable sampling without compromising the RBC unit. However, previous efforts to assess the RBC unit by sampling segments have relied on hemolysis and hemoglobin assays, which are not consistent in segments and blood bags^35,46,47^. Specifically, Kurach and colleagues previously investigated the potential to use blood unit segments for quality testing of blood bag during storage^35^. At 42 days, they observed that MCH and MCHC, but not MCV, were significantly greater in segments than in blood bags, for segments prepared using the same buffy coat method as our study. We performed direct linear comparison of all three RBC parameters and found strong correlation between blood bag and segment for all three. We noted relatively poor correlation between hemolysis in segment and blood bag. These differences may result from hemolysis being sensitive to differences in the extracellular chemical environment, such as the different plastics used to make the blood bag and segments, whereas changes in cell deformability are thought to result from intracellular proteomic remodeling and changes in oxidation^48^. Together, these data support the assessment of deformability, MCH, MCHC, and MCV on RBCs sampled from blood unit segments in place of blood bags in studies of blood transfusion efficacy.

Current practices in transfusion medicine dictate storing RBC units for up to 42 days, as well as using these units on a first-in-first out basis. However, such standards do not account for differences between donors, and specifically, differences in storage-based degradation. Determining these differences is particularly important for chronic transfusion recipients, since the storage-based degradation can reduce the circulation time of transfused RBCs, and thereby increase the frequency of transfusions for these recipients^2^. The ability to perform interim testing to RBC units during storage, without compromising the blood product, could therefore have a major impact on selecting RBC units for chronic transfusion recipients.

## Data availability statement

The data that support the findings of this study are available on request from the corresponding author.

## Ethics approval statement

This study was approved by the University of British Columbia’s Clinical Research Ethics Board (UBC REB# H19-01121) and Canadian Blood Services Research Ethics Board (CBS REB# 2019-029).

## Patient consent statement

Patient consent was not mandated for this study.

## Clinical Trial Registration

This study was not a clinical trial.

## Acknowledgments

We are grateful to Canadian Blood Services’ blood donors who made this research possible. We would also like to thank Dana Devine and Brankica Culibrk for helpful discussion and for providing access and training for the Sysmex system.

## REFERENCES

1. Bennett-Guerrero, E. et al. Evolution of adverse changes in stored RBCs. PNAS 104, 17063–17068 (2007).

2. D’Alessandro, A., Liumbruno, G., Grazzini, G. & Zolla, L. Red blood cell storage: the story so far. Blood Transfus 8, 82–88 (2010).

3. Matthews, K. et al. Microfluidic deformability analysis of the red cell storage lesion. Journal of Biomechanics 48, 4065–4072 (2015).

4. Tinmouth, A., Fergusson, D., Yee, I. C. & Hébert, P. C. Clinical consequences of red cell storage in the critically ill. Transfusion 46, 2014–2027 (2006).

5. Yoshida, T., Prudent, M. & D’Alessandro, A. Red blood cell storage lesion: causes and potential clinical consequences. Blood Transfus 17, 27–52 (2019).

6. Shander, A., Cappellini, M. D. & Goodnough, L. T. Iron overload and toxicity: the hidden risk of multiple blood transfusions. Vox Sanguinis 97, 185–197 (2009).

7. Wagner, S. J. Transfusion-transmitted bacterial infection: risks, sources and interventions. Vox Sang. 86, 157–163 (2004).

8. Hendrickson, J. E. & Hillyer, C. D. Noninfectious serious hazards of transfusion. Anesth. Analg. 108, 759–769 (2009).

9. Bux, J. Transfusion-related acute lung injury (TRALI): a serious adverse event of blood transfusion. Vox Sanguinis 89, 1–10 (2005).

10. Ng, M. S. Y., Ng, A. S. Y., Chan, J., Tung, J.-P. & Fraser, J. F. Effects of packed red blood cell storage duration on post-transfusion clinical outcomes: a meta-analysis and systematic review. Intensive Care Med 41, 2087–2097 (2015).

11. Berezina, T. L. et al. Influence of Storage on Red Blood Cell Rheological Properties. Journal of Surgical Research 102, 6–12 (2002).

12. Card, R. T., Mohandas, N. & Mollison, P. L. Relationship of post-transfusion viability to deformability of stored red cells. British Journal of Haematology 53, 237–240 (1983).

13. Huruta, R. R. et al. Mechanical Properties of Stored Red Blood Cells Using Optical Tweezers. Blood 92, 2975–2977 (1998).

14. Nagaprasad, V. & Singh, M. Sequential analysis of the influence of blood storage on aggregation, deformability and shape parameters of erythrocytes. Clinical Hemorheology and Microcirculation 18, 273–284 (1998).

15. Deplaine, G. et al. The sensing of poorly deformable red blood cells by the human spleen can be mimicked in vitro. Blood 117, e88–e95 (2011).

16. Franco, R. S. Measurement of red cell lifespan and aging. Transfus Med Hemother 39, 302–307 (2012).

17. Cohen, R. M. et al. Red cell life span heterogeneity in hematologically normal people is sufficient to alter HbA1c. Blood 112, 4284–4291 (2008).

18. Cicha, I. et al. Gamma-ray-irradiated red blood cells stored in mannitol-adenine-phosphate medium: rheological evaluation and susceptibility to oxidative stress. Vox Sang. 79, 75–82 (2000).

19. Suzuki, Y., Seiyama, A., Tateishi, N., Yamanishi, S. & Maeda, N. Rheological evaluation of X-ray irradiated blood. Vox Sang. 64, 139–144 (1993).

20. Suzuki, Y. et al. Decreased deformability of the X-ray-irradiated red blood cells stored in mannitol-adenine-phosphate medium. Clin. Hemorheol. Microcirc. 22, 131–141 (2000).

21. Laczkó, J., Feó, C. J. & Phillips, W. Discocyte--echinocyte reversibility in blood stored in CPD over a period of 56 days. Transfusion 19, 379–388 (1979).

22. Chien, S., Usami, S. & Bertles, J. F. Abnormal Rheology of Oxygenated Blood in Sickle Cell Anemia. J Clin Invest 49, 623- (1970).

23. Clark, M. R., Mohandas, N. & Shohet, S. B. Deformability of Oxygenated Irreversibly Sickled Cells. J Clin Invest 65, 189–196 (1980).

24. Clark, M. R., Mohandas, N., Shohet, S. B., Hoesch, R. M. & Rossi, M. E. Osmotic Gradient Ektacytometry - Comprehensive Characterization of Red-Cell Volume and Surface Maintenance. Blood 61, 899–910 (1983).

25. Lacelle, P. L. Alteration of Membrane Deformability in Hemolytic Anemias. Semin Hematol 7, 355- (1970).

26. Lacelle, P. L. Pathologic Erythrocytes in Capillary Microcirculation. Blood Cells 1, 269–284 (1975).

27. Lessin, L. S., Kurantsinmills, J. & Weems, H. B. Deformability of Normal and Sickle Erythrocytes in a Pressure-Flow Filtration System. Blood Cells 3, 241–262 (1977).

28. Snyder, L. M. et al. The Role of Membrane-Protein Sulfhydryl-Groups in Hydrogen Peroxide-Mediated Membrane Damage in Human-Erythrocytes. Biochim Biophys Acta 937, 229–240 (1988).

29. Snyder, L. M. et al. Effect of Hydrogen-Peroxide Exposure on Normal Human-Erythrocyte Deformability, Morphology, Surface Characteristics, and Spectrin-Hemoglobin Cross-Linking. J Clin Invest 76, 1971–1977 (1985).

30. Weed, R. I. Importance of Erythrocyte Deformability. Am J Med 49, 147- (1970).

31. Koch, C. G. et al. Duration of red-cell storage and complications after cardiac surgery. New Engl J Med 358, 1229–1239 (2008).

32. Högman, C. F. & Meryman, H. T. Red blood cells intended for transfusion: quality criteria revisited. Transfusion 46, 137–142 (2006).

33. Pavenski, K., Saidenberg, E., Lavoie, M., Tokessy, M. & Branch, D. R. Red Blood Cell Storage Lesions and Related Transfusion Issues: A Canadian Blood Services Research and Development Symposium. Transfusion Medicine Reviews 26, 68–84 (2012).

34. Islamzada, E. et al. Deformability based sorting of stored red blood cells reveals donor-dependent aging curves. Lab Chip 20, 226–235 (2020).

35. Kurach, J., Hansen, A., Turner, T., Jenkins, C. & Acker, J. Segments from red blood cell units should not be used for quality testing. Transfusion n/a-n/a (2013) doi:10.1111/trf.12303.

36. Guo, Q. et al. Deformability based sorting of red blood cells improves diagnostic sensitivity for malaria caused by Plasmodium falciparum. Lab Chip 16, 645–654 (2016).

37. Guo, Q., Duffy, S. P., Matthews, K., Islamzada, E. & Ma, H. Deformability based Cell Sorting using Microfluidic Ratchets Enabling Phenotypic Separation of Leukocytes Directly from Whole Blood. Scientific Reports 7, 6627 (2017).

38. Dumont, L. J. & AuBuchon, J. P. Evaluation of proposed FDA criteria for the evaluation of radiolabeled red cell recovery trials. Transfusion 48, 1053–1060 (2008).

39. Bratosin, D., Estaquier, J., Ameisen, J. c. & Montreuil, J. Molecular and Cellular Mechanisms of Erythrocyte Programmed Cell Death: Impact on blood Transfusion. Vox Sanguinis 83, 307 (2002).

40. Dhabangi, A. et al. Effect of Transfusion of Red Blood Cells With Longer vs Shorter Storage Duration on Elevated Blood Lactate Levels in Children With Severe Anemia: The TOTAL Randomized Clinical Trial. JAMA 314, 2514 (2015).

41. Fergusson, D. A. et al. Effect of fresh red blood cell transfusions on clinical outcomes in premature, very low-birth-weight infants: the ARIPI randomized trial. JAMA 308, 1443–1451 (2012).

42. Heddle, N. M. et al. Effect of Short-Term vs. Long-Term Blood Storage on Mortality after Transfusion. N Engl J Med 375, 1937–1945 (2016).

43. Lacroix, J. et al. Age of Transfused Blood in Critically Ill Adults. N Engl J Med 372, 1410–1418 (2015).

44. Steiner, M. E. et al. Effects of Red-Cell Storage Duration on Patients Undergoing Cardiac Surgery. N Engl J Med 372, 1419–1429 (2015).

45. Barshtein, G. et al. Deformability of transfused red blood cells is a potent determinant of transfusion-induced change in recipient’s blood flow. Microcirculation 23, 479–486 (2016).

46. Sowemimo-Coker, S. Red blood cell hemolysis during processing. Transfusion medicine reviews (2002).

47. Gyongyossy-Issa, M. I. C., Weiss, S. L., Sowemimo-Coker, S. O., Garcez, R. B. & Devine, D. V. Prestorage leukoreduction and low-temperature filtration reduce hemolysis of stored red cell concentrates. Transfusion 45, 90–96 (2005).

48. Kim-Shapiro, D. B., Lee, J. & Gladwin, M. T. Storage lesion: role of red blood cell breakdown. Transfusion 51, 844–851 (2011).

